# CTCF regulates anxiety and depression like behavior and maintenance of neuronal identity in the adult mouse brain

**DOI:** 10.1101/2023.11.02.565255

**Authors:** Liron Davis, Ipsita Chatterjee, Dmitriy Getselter, Evan Elliott

## Abstract

CCCTC-binding factor (CTCF) is a chromatin binding factor that binds to DNA sequence specific sites and, together with cohesin complex, establishes chromatin loops and regulates gene expression. CTCF was previously implicated as a major contributor in neural development. Genetic aberrations in CTCF are associated with intellectual disability, aggression, attention deficit, and autistic behavior. Previous mice-model studies have identified a necessary role for CTCF during development of CaMKIIa expressing excitatory neurons in the ability of learning and memory. However, it is not clear if CTCF is only necessary for development in the brain, or also in maintenance of neuronal functions and behavior in the adulthood. In the current study, adulthood-specific knockout of CTCF in excitatory neurons induced an elevation in anxiety and depression related behavior and a decrease in seeking social novelty. Depression and apathy-like behavior was reversed by treatment with serotonin specific reuptake inhibitor sertraline. Golgi staining analysis reveals major regression of dendritic complexity in the hippocampus and prefrontal cortex. In parallel, there is increased DNA compaction and decreased global H3K9 acetylation. Single nuclei RNA sequencing confirms a retreat in neuronal subtype identity in excitatory neurons after knockout. Gene ontology analysis display upregulation of genes that related regulation of cell population, neuronal development and neuronal differentiation and migration. These findings determine that CTCF is required for a propriate function of the mature excitatory neurons, independent of roles during development.

**Significance Statement:** The gene CTCF is an important regulator of gene expression. Mutations in CTCF have been identified in individuals with a range of neurodevelopmental disorders, including intellectual disability and autism. However, it is unknown if CTCF is important only for neuronal development, or plays functional roles in the brain during adulthood. The current study finds that CTCF deletion during adulthood in excitatory neurons induces a behavioral phenotype that that includes increase in anxiety and depression-like behavior and changes in social behavior. In addition, CTCF depletion in adulthood affects the morphology, identity and gene expression of these specific types of neurons. Therefore, CTCF is not only important in brain development, but also in maintenance of proper neuronal function and behavior during adulthood.

## Introduction

Genome organization in the nucleus of a eukaryotic cell is highly complex and dynamic, which includes chromosome territories, compartments and topologically associating domains (TADs) [1], [2]. CTCF, a zinc finger protein, has a pivotal role in organizing the genome within the nucleus and demarcates between the structural units of the chromosome [3]–[5]. CTCF works in tandem with cohesin complex, by making DNA physical contacts that create large loop domains, which regulate not only the 3D nuclear organization, but also gene expression including promotor-enhancer/ promotor-repressor interaction and RNA splicing.

Previous studies have reported that mutations in CTCF gene or CTCF binding sites are involved in neurodevelopmental and neurological diseases. Individuals with de novo mutations in CTCF gene developed intellectual disabilities, autistic features, impulsivity, microcephaly, and growth retardation [6]–[10]. The genetic mutations are located on different positions in the gene encoding CTCF (N-terminal region/ C-terminal region/ Zinc finger domain), and the clinical spectrum is highly variable. Moreover, modifications in CTCF binding sites have also been implicated in neuropsychiatric phenotype. For instance, CTCF directly binds to the promoter of FMR1 gene, the causative gene in Fragile × syndrome, and regulate its transcription [11].

Recently, several studies have shown that CTCF functions are dependent on developmental time point (embryonic stem cells/ mature cells). The influence of CTCF on the ability of embryonic stem cells (ESCs) to self-renew and differentiate is well established [1], [12]. In mice, CTCF deletion from oocytes results in embryo lethality by the morula stage, mediated by downregulation of p53 and upregulation of Puma [13], [14]. Studies shown that CTCF binds 40,000-80,000 sites genome wide and loss of CTCF results in 5000 differential gene expression after 96 hours of depletion in mouse ESCs [15], [16]. Therefore, CTCF has a critical role in cell proliferation, differentiation and survival by regulating the expression of multiple genes. The importance of CTCF not only in ESCs but also in cell maintenance of chromatin organization during postnatal and in the adult development.

A growing body of research in mouse models determined specific roles for CTCF in neuronal development and cognition. Depletion of CTCF from postmitotic neurons leads to lethality several weeks after birth [17], [18]. Furthermore, mice with a cKO of CTCF in postmitotic neurons had defects in dendritic arborization and synapse formation which include decrease in dendritic intersection, length and spine densityCTCF is particularly important for the proper neuronal expression of Pcdhs genes, which are essential for building functional neural networks in the brain (also neural diversity/identity). Notably, loss of γ-Pcdhs results reduced dendritic arborization [19]. Moreover, CTCF deletion from postnatal excitatory neurons displayed specific deficits in learning and memory, including spatial memory and fear memory [20], [21]. Taken together, these observations suggest that CTCF is required for survival and normal function of mature neurons by chromatin organization and gene expression.

However, all previous studies have deciphered the role of CTCF in organism development, either in developing neurons or in postmitotic neurons in early developmental periods. However, it is unknown if there is a role for CTCF in maintenance of proper behavior, neuronal morphology and identity, and gene expression during adulthood, after full organism development. In this study we investigate the role of CTCF in excitatory neurons by crossing mice carrying a loxP allele of CTCF [22] with mice transgenic for CaMKIIa-Cre-ERT, which induce conditional knockout (cKO) of CTCF in the CaMKIIa excitatory neurons. Treatment of mice with tamoxifen at eight weeks of age induces an adulthood excitatory forebrain neuron knockout. Adult mice with deletion of CTCF in CaMKIIa-expressing neurons exhibit increased anxiety and depression-like behavior. CTCF cKO mice displayed major changes in the dendritic morphology. In parallel, there is increased DNA compaction and decreased global H3K9 acetylation. In line with these finding, Single nuclei sequencing confirms a regression in neuronal identity and elevation in the number of gene which were downregulation of excitatory neurons after knockout.

## Materials and methods

### Mice

Mice were housed in a temperature-controlled barrier facility maintained at 24°C in a 12-hr light/dark cycle. Mice had access to fresh food and water ad libitum. To generate CTCF cKO specifically in forebrain excitatory neurons, CaMKIIa-Cre-ERT line was crossed to 𝐶𝑇𝐶𝐹^𝑓𝑙𝑜𝑥/𝑓𝑙𝑜𝑥^ mice. The double transgenic CaMKIIa − Cre −ERT^+/−^, 𝐶𝑇𝐶𝐹^𝑓𝑙𝑜𝑥/+^ were backcrossed to 𝐶𝑇𝐶𝐹^𝑓𝑙𝑜𝑥/𝑓𝑙𝑜𝑥^ mice to produce CaMKIIa − Cre − 𝐸𝑅𝑇2^+/−^,𝐶𝑇𝐶𝐹^𝑓𝑙𝑜𝑥/𝑓𝑙𝑜𝑥^ (cKO) and CaMKIIa − Cre −𝐸𝑅𝑇2^−/−^, 𝐶𝑇𝐶𝐹^𝑓𝑙𝑜𝑥/𝑓𝑙𝑜𝑥^ (WT) mice. These two lines were crossed in all experiments. Therefore, all offspring were 𝐶𝑇𝐶𝐹^𝑓𝑙𝑜𝑥/𝑓𝑙𝑜𝑥^, and half were with CaMKIIa − Cre − 𝐸𝑅𝑇2^+/−^. The mice without CaMKIIa − Cre − 𝐸𝑅𝑇2^+/−^, were the control animals. All experimental protocols were approved by the Animal Care and Use Committee of faculty of medicine, Bar-Ilan University, Israel.

To induce the expression of Cre in the Cre-ERT system, CTCF-cKO and WT control mice were treated with Tamoxifen at the age of 8 weeks. Tamoxifen was given once a day for four days (100 mg/kg per day; oral gavage) and the experiments started four weeks after the treatment.

### Sertraline treatment

Sertraline hydrochloride was purchased from Sigma-Aldrich. Sertraline was dissolved in a vehicle solution of 1% dimethyl sulfoxide (DMSO) in water. WT and CTCF-cKO mice were treated with vehicle (1% of DMSO in water) or sertraline (10 mg/kg) which administered i.p. 30 min before the behavioral tests.

### Behavioral testing

Mice were habituated to the behavioral room for at least 1 h before commencement of each test. Each test was performed on a separate day, with one day rest between each test. A camera filmed the movement, and the Noldus Software “EthoVision” was used to track the behavior of the animals.

### Rotarod test

The test was carried out with an accelerating Rotarod (Med Associates, St. Albins, VT). The speed of the Rotarod was set to 40 r.p.m. The amount of time each mouse spent on the rod was measured. The latency to fall was recorded three times with a 300 s cutoff time and average was calculated between the three independent trials.

### Wire test

The mouse was allowed to grip with their 2 forepaws a 2-mm thick horizontal metal wire and the latency to fall was recorded for two times, and the longest time until they full was recorded.

### Beam walk test

The mouse is placed on 1m horizontal beam with two different diameters (6mm, 12mm) and the time to reach an enclosed safety platform was measured. Before the test, mice were trained to traverse a beam for two days, twice each day.

### Open field test

Anxiety-like and locomotor behavior was determined in the open field. The size of the box is 50 x 50cm. The mouse is placed in a corner of the box and is allowed to explore for ten minutes. We recorded, using the EthoVision XT 10/11 software (Noldus), the distance moved and time spent in the entire box and in center square (25 x 25 cm). Testing was performed under light of 120 lux.

### Light/dark transition test

The mouse is placed in a dark plastic chamber (75X75 cm) with an opening to highly lit chamber (∼1200LUX). The mouse is free to move between the two chambers for five minutes. During this time, a camera films and tracks the behavior of the animals, including where they are found inside the box, velocity, distance traveled, etc.

### Elevated plus maze

The mouse is placed in the center of a four arms maze. Each arm is 30 cm in length, and two are closed and two are open. The maze is one meter high. The mouse is free to choose which arm it enters for a five minute period. During this time, a camera films and tracks the behavior of the animals, including where they are found in the maze, velocity, distance traveled, etc.

### Contextual and cued fear conditioning

First, on the training day each mouse was habituated to the fear conditioning cage for five minutes. Each mouse was placed into the conditioning chamber (10.5 × 10.5 × 10.5 cm) and allowed to explore freely for two minutes and a tone (75 dB) was sounded as the conditioned stimulus for 30 seconds followed by a two second mild foot-shock (0.7 mA) as the unconditioned stimulus. After a one-minute break, another tone and shock was administered and then the mouse was returned to the home cage one minute after the second tone-shock pair. The next day, the mice were placed back into the conditioning chamber for five minutes and their freezing behavior was measured during this time period as a measure of contextual memory. Three hours after context testing, the mice were placed into a different chamber with a novel odor, flooring, and light, for cue dependent memory testing. Following a two-minute habituation, the tone was presented thrice for 30 seconds with an interval of one minute in between each tone. Freezing during the three tone periods was recorded. The EthoVision XT 10-Noldus was used to analyze the videos. 3.5 Statistical analysis: Statistical Analysis Data were judged, and reported in figures and the figure legends, to be statistically significant when p < 0.05 by two-tailed t-test. Data are presented as mean ± SEM, and the number of animals (n) is mentioned.

### Y-maze test

Mice were placed at the junction between the arms of a symmetric Y maze. The mice allowed to freely explore the apparatus for 5 min. During this time, a camera films and tracks the continuity of arm entries. Spontaneous alternation was measured by counting the number of times the mouse entered each of the 3 arms of the maze in succession divided by the total number of alternations

### Social interaction test

The test took place in a Non-Glare Perspex box (60X40 cm) with two partitions that divide the box to three chambers, left, center and right (20X40 cm). The mouse is placed in the middle chamber for habituation (5 min) when the entry for both side chambers is barred. Test mouse was then allowed to explore the whole arena (10 min), where they freely choose between interacting with a novel mouse in one chamber or interacting with a familiar mouse in the other chamber. During this time, a camera films and tracks the behavior of the animals, including time spent in each chamber.

### Forced swim test

The mouse is placed in a dark plastic box, filled with water at 25C°to a depth of 15 cm for six minutes. Water was changed before the next animal was placed into the water tank. A camera films and tracks the duration of immobility during the test.

### Tail suspension test

The mouse is suspended by its tail 30 cm above the floor in a visually isolated area by adhesive tape that was placed ∼1 cm from the tip of the tail. Over six minutes of test, a camera films and tracks the duration of immobility.

### Sucrose preference test

was used to assess anhedonic behavior in mice. The test was carried out in home cages, while mice were single-housed by dividers. Each of the mouse was presented with two pipettes to choose freely: one containing drinking water and the other 0.5% sucrose solution. The test lasted 5 days, with the 1st day considered as habituation, while the next 4 days were considered as the test. During that time consumption of water and sucrose solution was measured.

### Immunostaining

Mice were perfused with 4% paraformaldehyde and then brains were dissected and incubated in 30% sucrose for two days. 30μm slices sections were taken by sliding microtome. Slices were blocked for one hour in blocking solution (10% horse serum, 0.3%triton and 1XPBS, and then incubated with primary antibodies for NeuN (1:500, Millipore) and CTCF (1:200, Millipore) or H3K9ac (1:400, Cell Signaling Technology) overnight at room temperature. The following day, slices were washed with incubated for 1 hour with secondary antibodies (alexa488 and cy3), stained for five minutes with Dapi (1:1000, Sigma), and washed three times, followed by mounting.

To visualize and evaluate chromatin condensation and total H3K9ac, the Stimulated emission depletion (STED) microscopy was applied. Image stacks were acquired at the sampling density of 80 nm per pixel and Z-spacing of 200 nm, using Plan Apochromat 63/1.4 Oil differential interference contrast (DIC) objective. To reduce noise and improve resolution, the stacks were 3D deconvolved by means of Huygens Professional software. Signal intensity quantification in individual nuclei (32 nuclei per group) was performed using Imaris software.

### Tunel staining

30 μm thick floating sections of 4% paraformaldehyde fixed mice brain were used to analyze apoptotic cells. The brain slices were processed using the In Situ Cell Death Detection Kits (Roche Life Science) according to manufacturer’s instruction.

### Golgi Staining

Performed on 12 weeks old mice, using the Golgi-Cox Stain kit (Bioenno Tech, LLC) according to the manufacturer’s procedures. Freshly dissected brain was sectioned at 150 mm on a microtome and mounted. For the neuronal morphology analysis, pyramidal neurons and granule cells were randomly selected from prefrontal cortex layer III/IV, hippocampal CA1 and DG. Images taken using bright field microscopy, and dendritic length and sholl analysis were analyzed using Imaris 9.1.2 software. Statistical analyses were performed using the two-way anova with GraphPad Prism 5 software.

### Corticosterone and Stress-Induction Protocol

Blood samples collected in four time points. Basal corticosterone levels were collected before the acute stress procedure. Stress-induced corticosterone levels were obtained from animals (n = 8 per group) that were restrained in 50-ml tubes for 30 min. Blood samples collected immediately after the acute stress, 30 min and 90 min after the acute stress. Blood was collected in tubes containing 0.5M EDTA. The blood was centrifuged at 2,000rcf for 30 min in 4°C, and serum was collected and frozen at −80°C until the assay was performed. Quantification of plasma corticosterone levels was carried out using the corticosterone RIA kit (MP Biomedicals, LLC) according to the manufacturer’s protocol.

### Single-nucleus extraction, RNA sequencing, and analysis

For the single-nucleus RNA-seq experiment, two weeks old mice were sacrificed. The forebrain of each mouse were dissected and minced using homogenizer in 5 ml homogenization buffer (0.25M sucrose, 25mM KCl, 5mM MgCl2, 20mM Tricine-KOH, 1mM DTT, 0.15mM spermine, 0.5mM spermidine), 7 µl RNase Inhibitor (Promega N2611) and 300 µl 5% IGEPAL. The sample was filtered through a 40 µm strainer, mixed with 5 ml of 50% iodixanol (Sigma D1556), underlayed with a gradient of 30% and 40% iodixanol, and centrifuged at 4,600g for 1 hour in a swinging bucket centrifuge at 4°C. Nuclei were collected at the 30%-40% interface. The nuclei was transferred to a 2 ml tube and centrifuged 10 min at 1,000*g* and 4 °C. Cell pellet was resuspended in PBS 1×, 0.04% BSA. Libraries for single nuclei RNA-seq were prepared using the Chromium Next GEM Single Cell 3′ GEM, Library & Gel Bead Kit v3.1 as recommended by the manufacturer. Cell-RT mix was prepared to aim 10,000 nuclei per sample and applied to Chromium Controller for GEM generation and barcoding. Libraries were sequenced on a NextSeq 550 sequencing system to a depth of approximately 30,000-40,000 reads per cell. Raw sequencing reads were analyzed using 10X Genomics Cell Ranger version 4.0.0 pipeline. Following fastq generation, counting was performed using a custom “pre-mRNA” reference package (based on Mus musculus mm10 genome), listing each gene transcript locus as an exon, in order to count intronic reads. Cell loupe browser was used to perform differential gene analysis and to visualize data. Mitochondrial genes (genes beginning with mt-) were taken out of the final analysis, as they should not be present in nuclei, and are likely indicative of a technical artifact. Gene ontology analysis was performed using Toppgene.

### Statistical analysis

Statistical Analysis Data were judged, and reported in figures and the figure legends, to be statistically significant when p < 0.05 by two-tailed t-test. Data are presented as mean ± SEM, and the number of animals (n) is mentioned.

## Results

### Conditional knockout of CTCF in Excitatory adulthood neurons

In order to determine the effects of CTCF depletion in mature excitatory neurons on behavior and neuronal function, we used the CaMKIIa-Cre-ERT model. We crossed *CTCF*^loxP^ mice with CaMKIIa *-Cre- ERT* mice, which express Cre recombinase under the control of the forebrain excitatory neuron-specific promoter for CaMKIIa. To induce the Cre-mediated knockout of CTCF, cKO and WT control mice were treated with Tamoxifen at eight weeks of age.

Immunohistochemistry analysis validated knockout of CTCF in neurons of the hippocampus (Fig. 1A). Surprisingly, unlike other excitatory neuron CTCF knockout models, where the mice displayed a shortened lifespan, adulthood knockout animals do not display enhanced lethality. CTCF-cKO mice displayed no apparent health abnormalities, normal growth and survived over one year (which was the last age we checked) (Fig. 1B, C). Tunel staining analysis revealed no apoptosis in the hippocampus of knockout mice (Fig. 1D). In addition, CTCF-cKO mice did not exhibit any abnormal level of locomotion in rotarod test, wire hang test or beam test (Fig. 1E-G).

**Figure 1:**
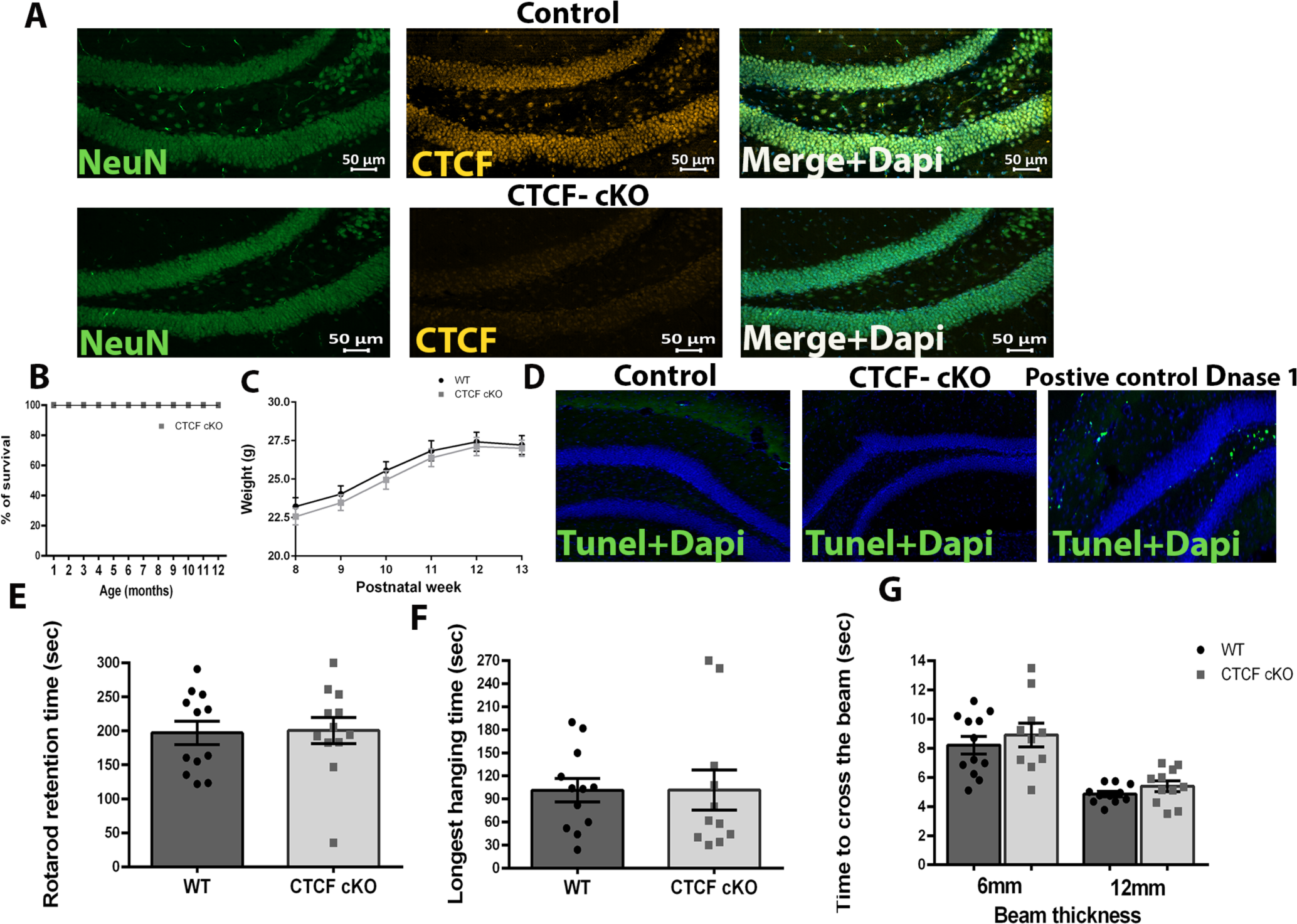
Characterization of the CaMKIIa-Cre-ERT- Mediated CTCF- cKO Mice. (A) Immunofluorescence analysis of CTCF in the hippocampus of twelve-week-old WT and CTCF- cKO mice shows depletion of CTCF from hippocampal neurons. (B) Lifespan (C) body weight and (D) tunel staining analysis of CTCF- cKO mice. No animals died over one year after birth and no significant changes in the weight of CTCF- cKO mice. CTCF- cKO, n = 17; WT, n = 22 mice per group. (E-G) CTCF- cKO mice displayed no deficit in motor skills. (E) CTCF- cKO mice spent similar time on the rotator, (F) on the wire, and (G) the time spent to cross the 6mm/ 12mm thickness beam was the same between WT and CTCF- cKO mice. CTCF- cKO, n = 12; WT, n = 12 mice per group, two-tailed t-test. Values in graphs are expressed as the mean ± SEM.

### Anxiogenic related behavior and dysfunction in sociability in CTCF- cKO mice

To evaluate effects of adulthood knockout on behavior, a battery of behavioral tests were performed four weeks after tamoxifen treatment. First, we analyzed anxiety-like behavior. In open field test, CTCF- cKO mice spent significant less time in the center (Fig. 2A; * p=0.046) and lower number of entries into the center (Fig. 2B; ** p=0.0071). Although the mice display no dysfunction in motor skills, as shown above, the distance that the CTCF- cKO mice travelled in the open field environment was declined (Fig. 2C; ** p=0.0018). In dark light test CTCF- cKO mice spent significant less time in the light zone (Fig. 2D; * p=0.0155) and equal number of entries into the light chamber (Fig. 2E). In the elevated plus maze test CTCF-cKO mice spent less time in the open arms (Fig. 2F; ** p=0.0021) and had a smaller number of entries into the open arms (Fig. 2G; * p=0.0285). Thus, three separate tests indicate that knockdown of CTCF in mature excitatory neurons during adulthood heightened anxiety-related behavior.

**Figure 2:**
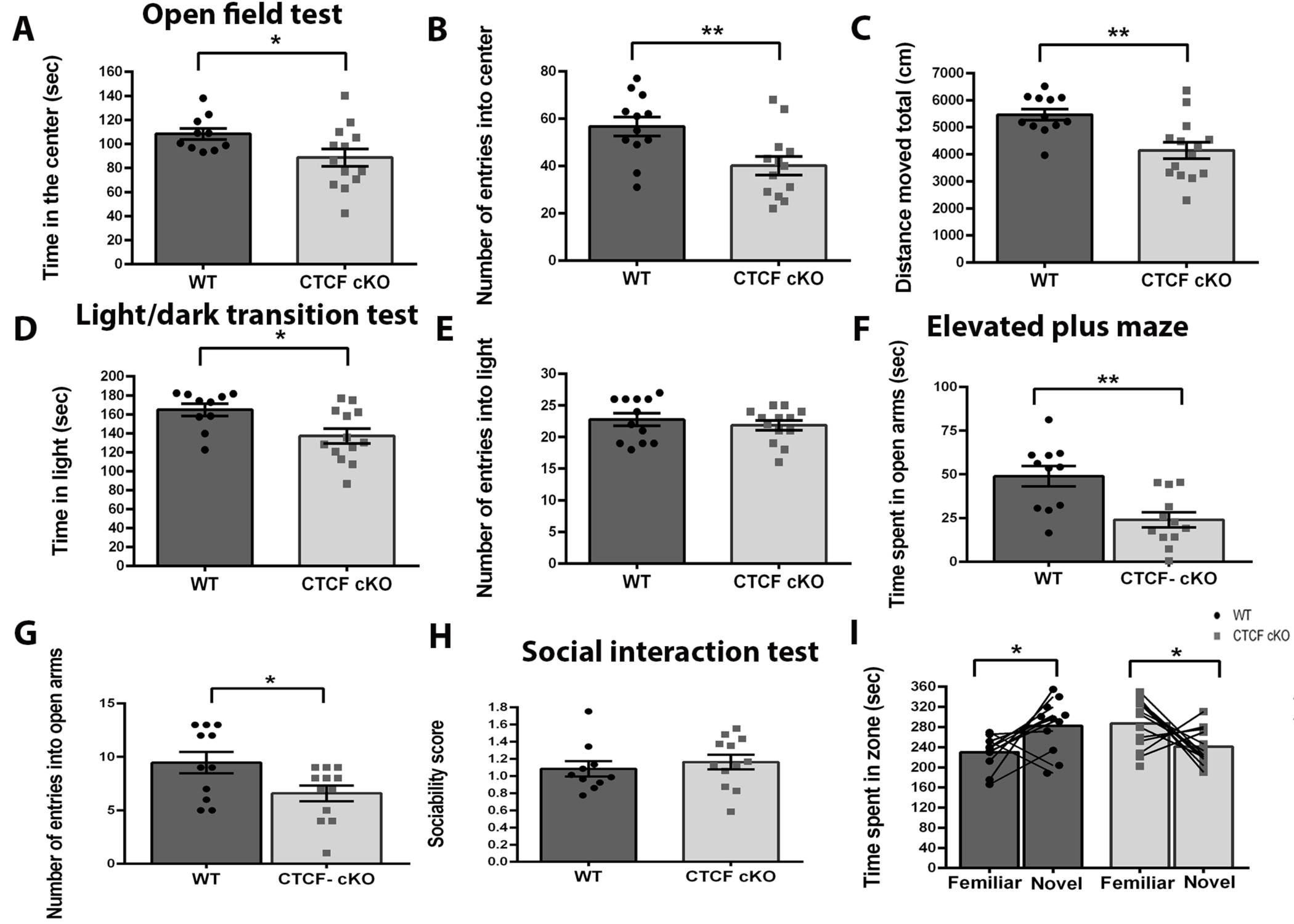
Heightened anxiety-related behavior and dysfunction in social recognition in CTCF- cKO mice. In the open field test, (A) CTCF cKO mice spent less time in center (*p=0.046) and (B) lower number of entries into center (**p=0.0071). (C) Distance traveled by CTCF- cKO mice in the open-field arena was lower compered to WT mice (**p=0.0018). In the light/dark transition test, (D) CTCF cKO mice spent less time in light zone (*p=0.0155), (E) with no differences in the number of entries into light zone. In the elevated plus maze test, (F) CTCF cKO mice spent less time in open arms (**p=0.0021) and (G) had lower number of entries into open arm (*p=0.0285). (H-I) Results of the three-chambered social test, (H) CTCF- cKO mice displayed no dysfunction in sociability. (I) However, unlike WT mice, CTCF- cKO mice prefer familiar mouse compared to a novel mouse in social recognition test (WT, *p=0.0126; cKO, *p=0.0149). CTCF- cKO, n = 13; WT, n = 11 mice per group; two-tailed t test. Values in graphs are expressed as the mean ± SEM.

In three chamber social interaction test, CTCF-cKO mice showed no defects in sociability as defined by preference for social interaction in comparison to empty chambers (Fig. 2H). However, in social novelty paradigm, the CTCF-cKO mice showed preference to be in the zone of the familiar mouse compared with the novel mouse, unlike WT mice (Fig. 2I; WT, *p=0.0126; cKO, *p=0.0149). This data suggests that knockdown of CTCF in CaMKIIa excitatory neurons changes social preference behavior.

### CTCF- cKO mice displayed depression like behavior with no changes in learning and memory

In order to examine the effect of adulthood CTCF deletion in excitatory mature neurons on depression-like behavior, we performed tail suspension test, forced swim test and sucrose preference tests. In both tail suspension and forced swim tests, CTCF- cKO mice spent more time immobile compared to the control group (Fig. 3A, B; ** p=0.0027, ** p=0.0011respectively). Furthermore, in the sucrose preference test, CTCF- cKO mice showed a significant reduction in their preference for sucrose over water compared to WT mice (Fig. 3C; *p=0.0479). Therefore, these three tests indicate that low expression of CTCF in mature excitatory neurons heightened depression like behavior.

**Figure 3:**
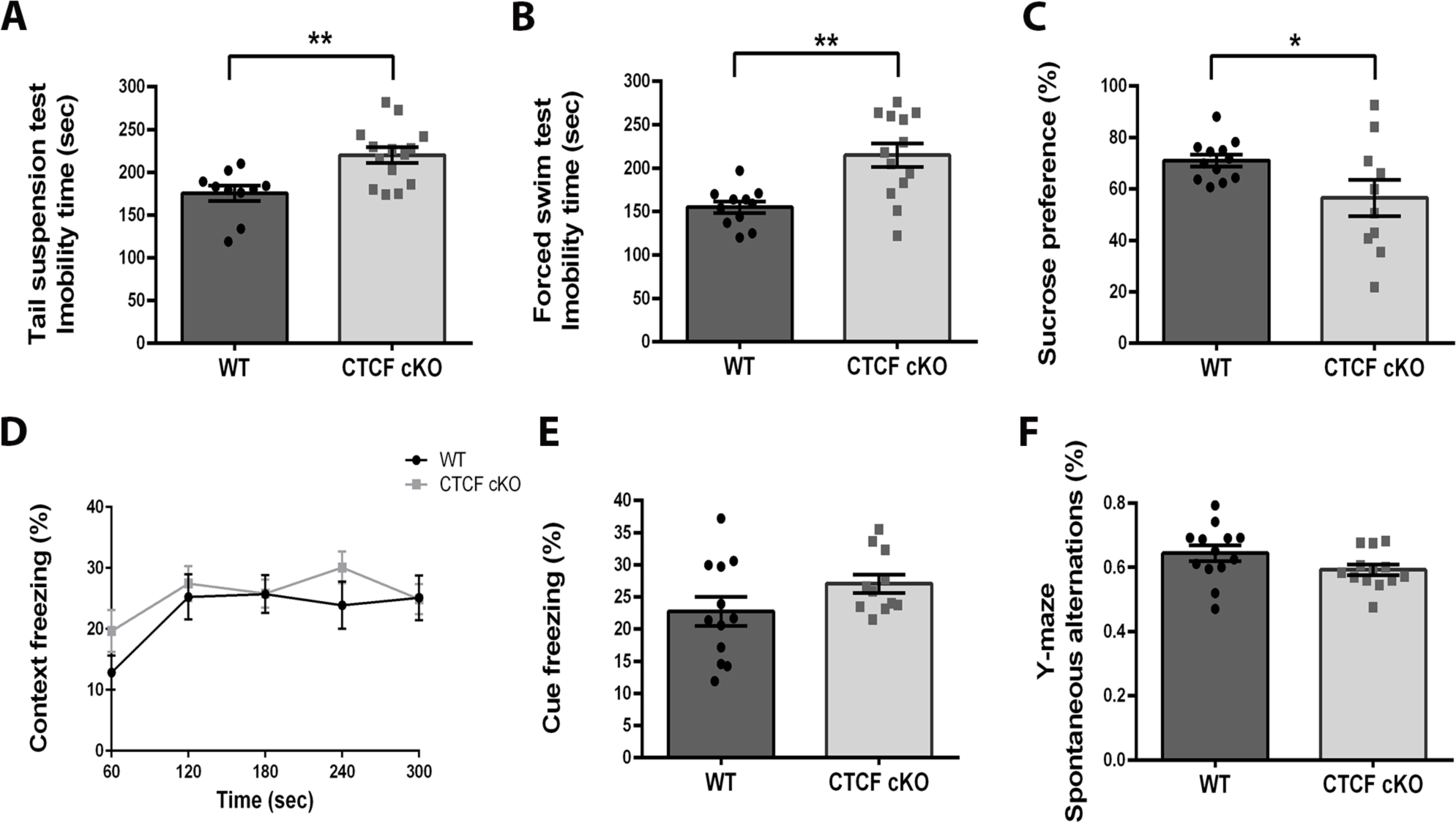
Heightened depression-related behavior with no changes in learning and memory in CTCF- cKO mice. Performance in depression behavioral tests, (A-B) CTCF cKO mice spent more time immobile compared to the WT group in (A) tail suspension test (**p=0.0027) and (B) forced swim test (**p=0.0011). In the sucrose preference test, (C) CTCF cKO mice showed a significant reduction in sucrose preference over water compared with WT mice (*p=0.0479). (D-F) Performance in learning and memory tests, (D-E) CTCF cKO mice showed proper freezing compered to WT group in the contextual and cue-dependent fear conditioning test. (F) In Y-maze test, CTCF cKO mice had similar spontaneous alternation compered to WT group. CTCF- cKO, n = 11; WT, n = 12 mice per group; two-tailed t test. Values in graphs are expressed as the mean ± SEM.

Since depletion of CTCF in excitatory neurons during developmental time point induced deficits in learning and memory in the fear extinction paradigm, we examined if deletion in the adulthood excitatory neurons cause the same phenotype. In contextual and cued fear conditioning test, CTCF- cKO mice displayed similar percentage of freezing levels (Fig. 3D, E). Further, in Y-maze test, CTCF cKO mice had proper spontaneous alternation compered to WT group (Fig. 3F). These results suggest that CTCF deletion in adulthood postmitotic excitatory neurons does not induce dysfunction in learning and memory.

### No differences in Blood corticosterone levels of CTCF- cKO mice compared to WT mice

Due to the anxiety-like phenotype, we hypothesized that these mice may display a dysfunction in the Hypothalamic-Pituitary-Adrenal (HPA) axis. We measured levels of blood corticosterone in both mice groups before and after acute stress. Blood was taken from mice tail in 4 time points; before the acute stress (0), and different time points (30min, 60 min, and 120min) after the initiation of stress. There were no significant differences in corticosterone levels between CTCF- cKO mice and WT littermates. (Fig. 4A). Therefore, differences in the HPA-axis cannot explain the anxiety-like phenotype in these mice.

**Figure 4:**
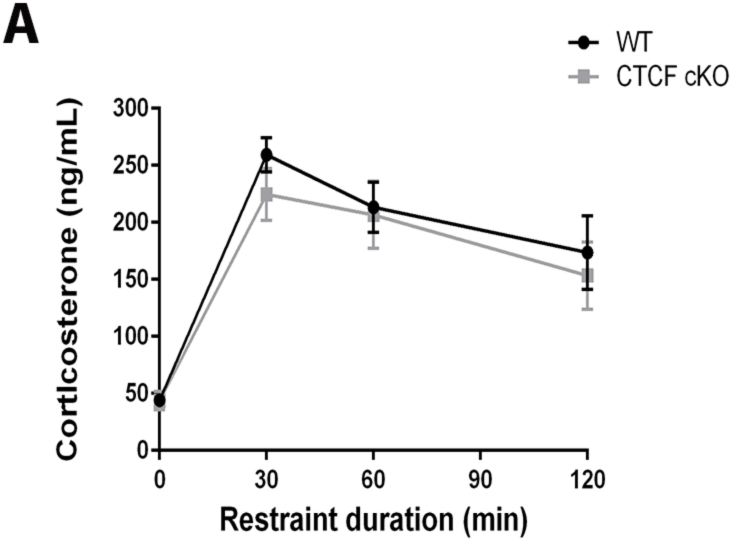
normal levels of corticosterone in CTCF- cKO mice. (A) RIA analysis of corticosterone levels of twelve weeks old CTCF- cKO and WT mice revealed no changes between the groups. n= 8 mice per group, two-tailed t-test. Values in graphs are expressed as the mean ± SEM.

### Rescue of depression-like behavior in adult CTCF- cKO mice by treatment with sertraline

In order to confirm depression-like behavior in CTCF- cKO mice, we studied the effect of antidepressants on depression-related and locomotion phenotypes. To evaluate the efficacy of sertraline treatment on behavioral abnormalities in CTCF- cKO mice, exploratory activity and depression like behavior tests were performed. Sertraline, a Serotonin reuptake inhibitor (SSRI), or saline vehicle, was injected i.p and behavioral tests were performed 30 min after acute treatment. Acute administration of sertraline attenuated the decreased exploratory behavior in open field in CTCF- cKO mice (Fig. 5A; ANOVA Tukey posthoc test). Furthermore, in open field test, the number of entries into center was similar between WT and CTCF- cKO group after administration of sertraline (Fig. 5B). In addition, sertraline attenuated the decrease in the duration of immobility in the CTCF- cKO mice in tail suspension test and a forced swimming test (Fig. 5C, D).

**Figure 5:**
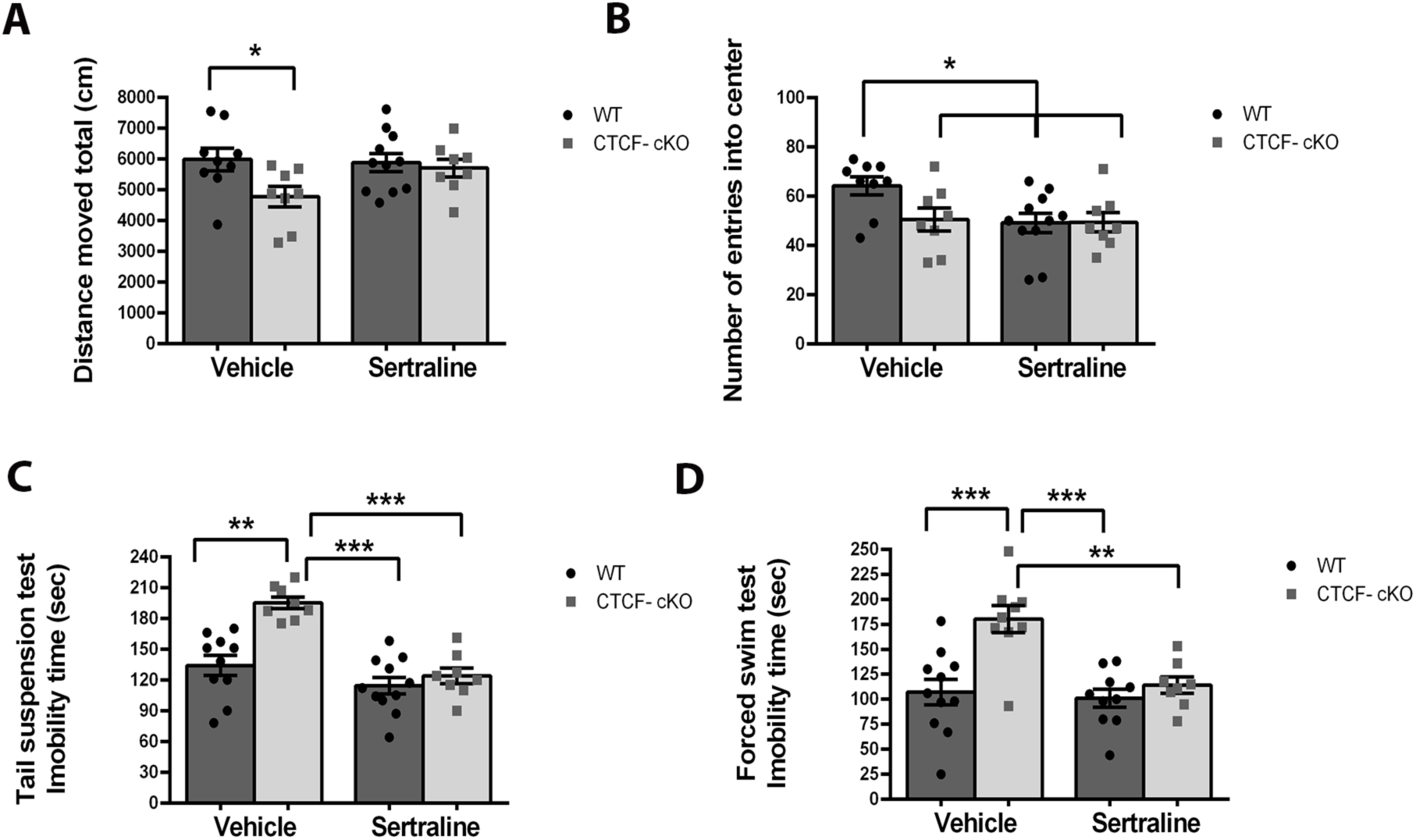
Treatment with sertraline rescues depression like behavior in CTCF- cKO mice. (A) Locomotor activity measured as total distance traveled during open field test in vehicle-treated and sertraline-treated, twelve weeks old mice. vehicle-treated CTCF- cKO mice traveled a significantly lower distance (*p =0.013) as compared to WT mice, while treatment with sertraline significantly increased locomotor activity in CTCF- cKO mice. (B) vehicle-treated CTCF- cKO mice had lower number of entries into center (*p =0.04) as compared to WT mice, while treatment with sertraline significantly increased the number of entries into center in CTCF- cKO mice. Furthermore, vehicle-treated WT mice had higher number of entries into center as compared to sertraline-treated WT mice (p=0.013), and CTCF- cKO mice (p=0.025). (C-D) Performance in depression behavioral tests, (C) vehicle-treated CTCF- cKO mice spent more time immobile compared to vehicle and sertraline-treated WT group (**p= 0.002, ***p<0.001 respectively), while treatment with sertraline significantly increased their mobility in tail suspension test (***p<0.001). (D) Similar result was display in forced swim test (***p<0.001, ***p<0.001, **p= 0.002, respectively). vehicle-treated: CTCF- cKO, n = 8; WT, n = 9 and sertraline-treated: CTCF- cKO, n = 8; WT, n = 11; two-way ANOVA. Values in graphs are expressed as the mean ± SEM.

These data suggest that decreased exploratory behavior and depression-like behavior in the CTCF- cKO mice was reversed and responsive to acute administration with antidepressants that selectively inhibit SERT, such as sertraline. This gives evidence that the decreased activity in open field is due to a depressive-like phenotype in these mice, and may be a form of apathy.

### CTCF- cKO mice display major reduction in the number of dendritic arborization

To examine the morphology of pyramidal neurons in the prefrontal cortex and hippocampal CA1 and DG in CTCF-cKO mice versus controls, we used Golgi-Cox staining in hippocampus of mice that were perfused four weeks after tamoxifen treatment. Sholl analysis showed that in pyramidal cells, in both hippocampal Ca1 and prefrontal cortex, there was significant reduction in apical and basal dendrite arborization in CTCF-cKO mice (Fig. 6A, B; ***p<0.001).

**Figure 6:**
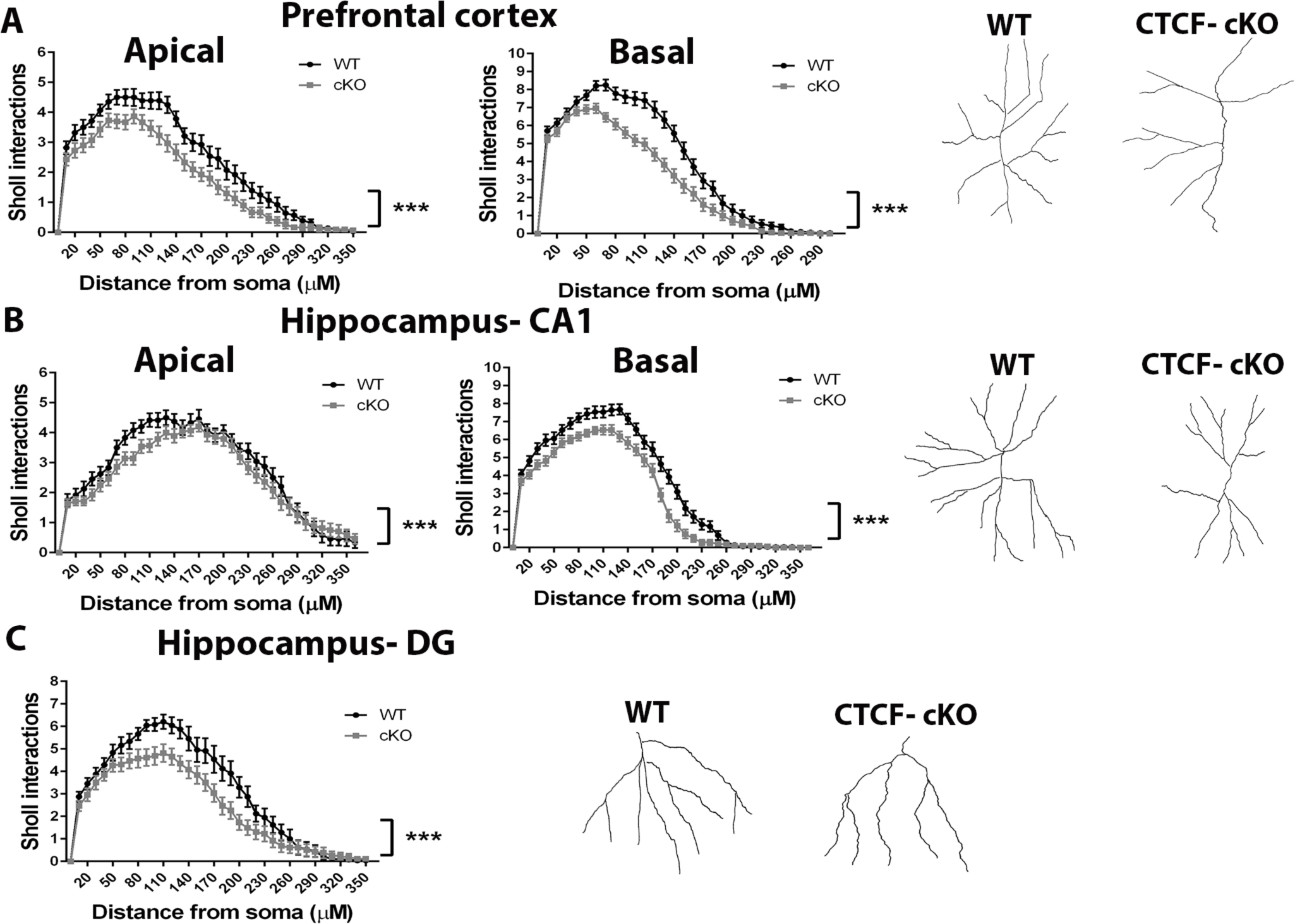
Golgi staining of prefrontal cortex and hippocampus of twelve weeks old CTCF- cKO mice determine major reduction in the number of dendritic arborization. Sholl analysis was used for examining dendritic branches in WT and CTCF- cKO mice. (A) Apical and Basal dendrites of CTCF- cKO mice are significantly less branched in the prefrontal cortex. In addition, (B) there is a decrease in dendritic intersection in the apical and basal dendrite of hipocample Ca1 and (C) dendrite of the dentate gyrus. n= 4 mice per group, two-way ANOVA. Values in graphs are expressed as the mean ± SEM. n = 28 neurons per group ***p<0.001

Additionally, in the granule cells of the DG, there was a decrease in dendritic branching (Fig. 6C; ***p<0.001). These data demonstrated that CTCF is important for the maintenance of dendrite morphology in neurons during adulthood.

### Aberrant chromatin architecture in hippocampal neurons in CTCF-cKO mice

CTCF, as DNA binding protein, is known for its essential role in chromatin architecture and genome organization. Therefore, we evaluated the effect of CTCF on chromatin architecture in hippocampal neurons. Quantitative 3D analysis of neuronal nucleus that were labelled with the DNA stain 4’,6-diamidino-2-phenylindole (DAPI) confirmed the increase in the number of intranuclear DAPI-stained foci in pyramidal neurons (Fig. 7A, B; *p=0.0160) and revealed that these structures were larger than in control mice (Fig. 7A, C; **p=0.0013). Furthermore, quantitative 3D analysis of global hypoacetylation of acetyl-histone (H3K9ac) confirmed the decrease in global H3K9ac in hippocampal neurons in CTCF-cKO mice compared to WT mice (Fig. 7A, D; ***p<0.001). These results suggest CTCF has a role in the maintenance of DNA compaction in timepoints after development.

**Figure 7:**
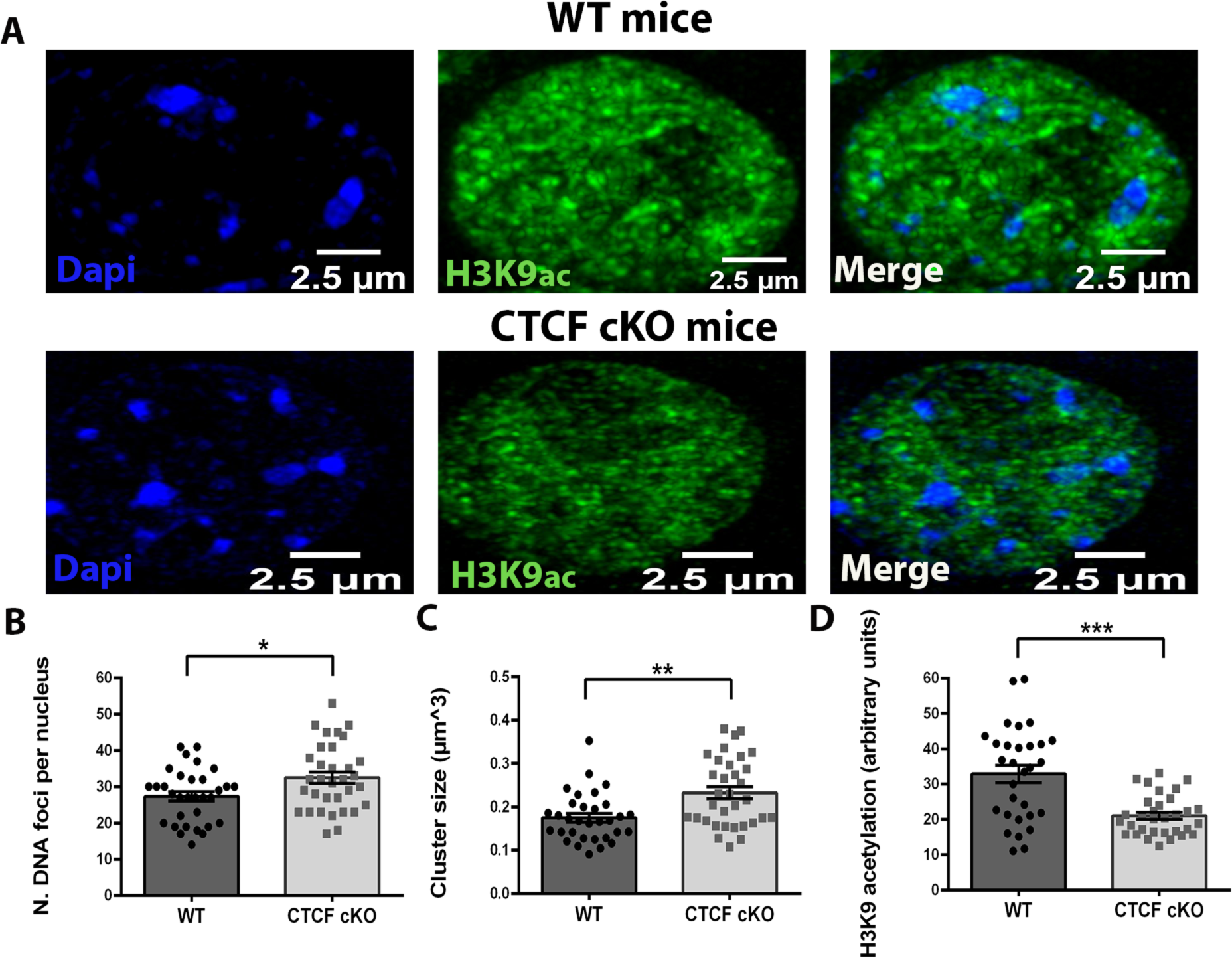
<u>Dysregulated chromatin in adulthood CTCF- cKO mice.</u>. (A) 3D reconstruction of confocal images illustrating both heterochromatin and H3K9 acetylation in hippocampal neurons of CTCF- cKO mice compered to WT. Quantification of the (B) number and (C) size of DAPI-stained foci per nucleus in CTCF- cKO mice demonstrate elevation compared to the control group (*p=0.0160, ** p=0.0013respectively). (D) Quantification of H3K9 acetylation display decrease in CTCF- cKO mice compared to the control group (***p<0.001). n= 4 mice per group, two-tailed t-test. Values in graphs are expressed as the mean ± SEM. n=32 nuclei per group.

### Single nuclei RNA sequencing confirms a regression in neuronal identity of excitatory neurons after depletion of CTCF

To understand the possible roles for CTCF in gene transcription and neuronal subtype identity, we performed single-nuclei RNA-seq (snRNA-seq) from hippocampus of CTCF- cKO and WT mice four weeks after tamoxifen treatment. Approximately 10,0000 nuclei were sequenced from each experimental group, and approximately 20% of those nuclei were Camkiia positive (Fig. 8A). Using known markers for specific excitatory neuron subtypes we were able to identify different subpopulations: Granule cells, Interneurons, CA1-pyramidal neuron A, CA1-pyramidal neuron B, CA3-pyramidal neuron (Fig. 8B, Fig. 8-1). Surprisingly, analysis that separated between WT and CTCF- cKO nuclei exposed that CTCF- cKO nuclei in all subpopulations are found closer into center in UMAP plot (Fig. 8C). This indicates a decrease in cell-specific identity, as has been previously reported [23].

**Figure 8:**
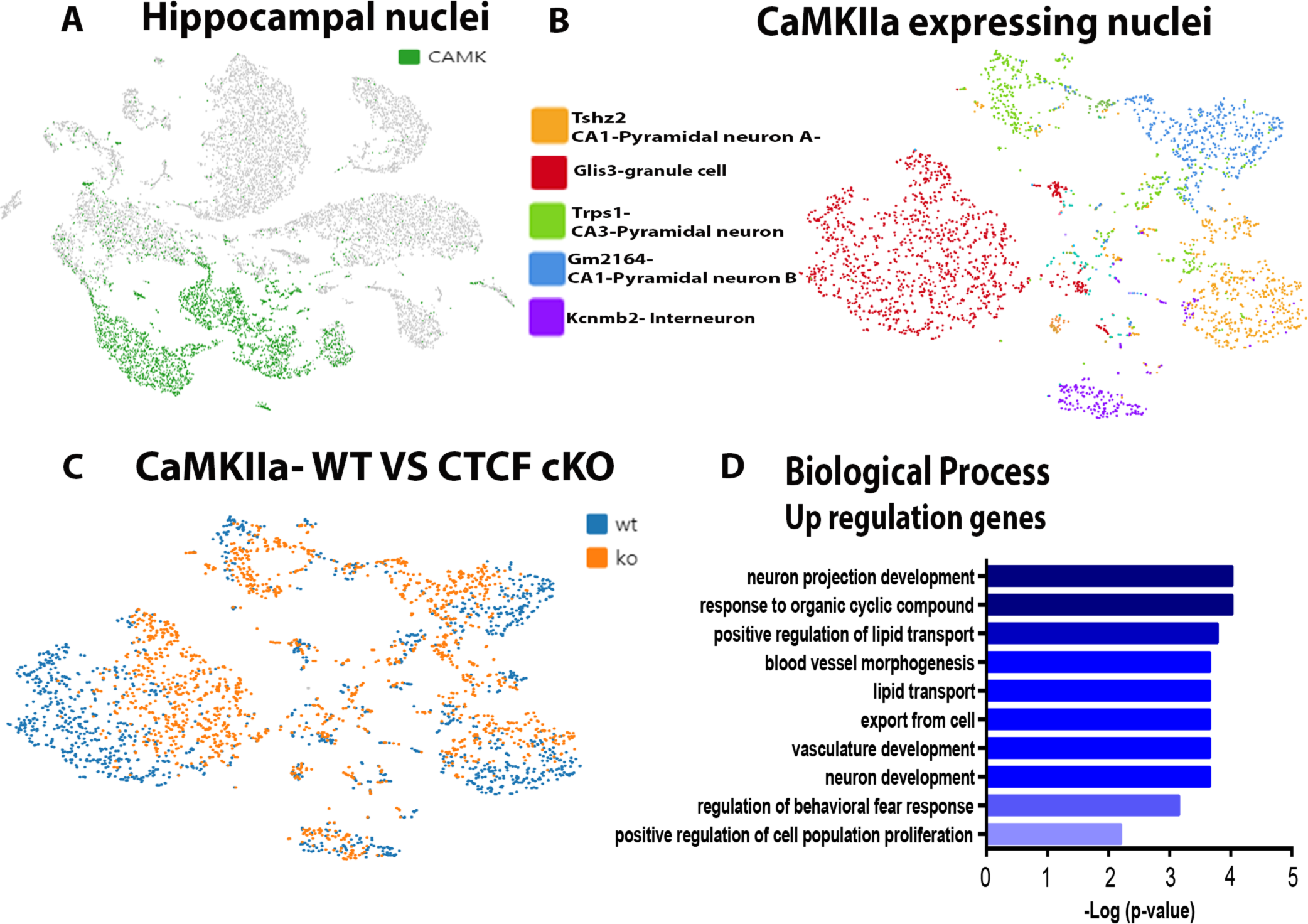
Single-nucleus RNA sequencing exposed a retreat in neuronal identity of excitatory neurons after CTCF conditional knockout. (A-B) UMAP plot of integrated analysis of snRNA-seq datasets from hippocampal neurons of CTCF- cKO and control littermates. (A) The cluster of the CaMKIIa expressing excitatory neurons marked in green. (B) Reclustering specifically of CaMKIIa expressing excitatory neurons. The identified subpopulations of the CaMKIIa expressing excitatory neurons: Granule cell, Interneuron, CA1- pyramidal neuron A, CA1- pyramidal neuron B, CA3- pyramidal neuron. Nuclei are colored by their classification label as indicated. (C) Single-nucleus analysis of hippocampal excitatory subpopulation neurons reveals cell state-transitions between CTCF- cKO (orange) compared to the control group (blue). (D) GO analysis of upregulation expressed genes highlighted genes that related to regulation of cell population, neuronal development and differentiation.

Furthermore, we performed differential expression analysis specifically on the CaMKIIa-expressing neurons and found 470 genes that were differentially expressed in the CTCF- cKO CaMKIIa- expressing nuclei compared to WT CaMKIIa- expressing nuclei (Supplementary Tables 1,2), including 176 upregulated genes and 294 downregulated genes. Gene ontology analysis revealed that upregulated genes were enriched for biological categories dealing with neuron development and cell proliferation (Fig. 8D). It is of interest to note an enrichment, as well, in genes involved in the behavioral fear response (such as CCK), which correlated with the anxiety phenotype. The findings of upregulation in development and proliferation related genes further suggest that the cells were reverting to a gene expression program of development, which correlates with the decreased dendritic complexity seen in the golgi staining and the retraction to center of the UMAP plot seen in the single cell sequencing. Therefore, the single cell sequencing data further suggests that CTCF is necessary for maintenance of a gene expression program relevant to maintaining a fully developed state in neurons.

## Discussion

The primary aim of this study is to understand the role of neuronal CTCF in mature CaMKIIa expressing excitatory neurons during adulthood. The main importance of this study lies in the attempt to understand if this genome organizer is also necessary for behavioral and neuronal maintenance during adulthood of the organism, or is only necessary for proper development.

The main behavioral phenotypes of selective knockout during adulthood in excitatory neurons is increased anxiety- and depression-like behavior, together with a deficit in social novelty behavior. While changes in anxiety-like behavior and social novelty have been reported in knockout of CTCF during developmental time periods, increase in depression-like behavior is a novel finding that may be related to a unique role of CTCF in adulthood. Some interesting previous findings support a specific role for CTCF in depression-like behaviors. First, several studies have found that CTCF regulates the expression of the serotonin transporter [24]–[26]. This may explain why SSRI treatment in our study was able to attenuate the depression-like phenotype.

Second, the increase of Kcnj8, which encodes for the protein Kir6.1, a potassium voltage-gated channel, seen in the single nuclei RNA-seq analysis may be a possible mechanism of the increase in depression behavior. Kir6.1 channel blockade by memantine improves depressive-like behaviors in mice [27]. Second, a recent study compared large cohort of schizophrenia and bipolar disorder brains. In the study, they mapped and found large clusters of hyper-acetylated regulatory domains, which strongly enriched for occupancies of the structural protein CTCF and were enriched for schizophrenia heritability[28].

In the present study of CTCF, we have found that knockdown of CTCF in mature CaMKIIa expressing excitatory neurons induces a decrease in seeking social novelty. As explained above, some patients with CTCF mutations are diagnosed with autism, which includes social deficits. There have been conflicting results in previous studies of CTCF knockdown in developmental excitatory neurons whether that knockdown produces direct effects on sociability [18]. Therefore, our results verified that CTCF may regulate social behavior partially through effects in excitatory neurons.

Some of the individuals with de novo mutation of CTCF observed temper tantrums and aggressivity. It well known that temper tantrums are often driven by anxiety in younger children. Furthermore, low expression of cohesion (conditional Smc3+/− knockout), which functions together with CTCF to establish chromatin loops, leads to heightened anxiety-related behavior in adult mice [29]. In our study, CTCF- cKO in adulthood neurons specifically increased anxiety like behavior. The lack of differences in corticosterone suggest that the anxiety changes aren’t due to dysregulation of the Hypothalamus-Pituitary-Adrenal Axis. The increase of CCK, an anxiogenic peptide, seen in the single nuclei RNA-seq analysis may be a possible mechanism of the increase in anxiety behavior.

Previous studies determined that developmental knockout of CTCF in excitatory neurons induce deficits in learning and memory with only subtle changes in neuronal morphology[21], [30], [31]. However, we have shown that knockdown of CTCF in mature excitatory neurons doesn’t cause learning and memory dysfunction. However, there was a very significant decrease in dendrite complexity in the hippocampus and prefrontal cortex with sociability impairment. These data suggest that CTCF has a different role in different time points during neuronal development. These results were surprising for us, because we assumed that CTCF is more critical in neuronal development than neuronal maintenance in the adulthood. These finding suggests a crucial role of CTCF specifically in maintaining correct neuronal morphology specifically in the adulthood stage.

In this study, we also provide evidence that CTCF affects DNA condensation and total H3K9 histone acetylation. CTCF can regulate transcriptional activation or repression. Both mice and human RNA-seq based studies show that CTCF contributes more towards gene activation than gene inactivation in the hippocampus, cortical cultures and in lymphocytes[9], [21], [30]. Furthermore, there are several studies that determined an overlap between CTCF enrichment and histone modification (including H3K9ac) on the genome [28], [32]–[34]. Therefore, this may explain the decrease in H3K9ac upon CTCF deletion. Moreover, the common assumption is that loss of CTCF does not impact A and B genomic compartmentalization, but concomitantly disrupted loops between TAD boundaries and insulation of neighboring TADs [35]. However, recent study of Zhang et al, claim that the role for CTCF in chromatin compartmentalization previously was underappreciated [36]. This study suggests that removal of CTCF may allow cohesin to travel beyond CTCF-binding sites, thereby increasing loop sizes. Structural loops originating within A-type compartment domains may thus extend into flanking B-type compartment domains and increase their contact probability. Increased contact probability can theoretically lead to increased condensation. Therefore, these studies are compatible with our results that present the influence of CTCF on DNA compaction and overall histone acetylation.

At the transcriptional level, CTCF- cKO CaMKIIa expressing excitatory nuclei displayed more downregulation genes (294) then upregulation genes (186). The upregulated genes were involved in biological process such as regulation of cell population, neuron development and neuron differentiation. The changes in gene expression, particular in genes that related neuronal development, also with the dendrite retreat we reveals in the Golgi staining emphasize the role of CTCF in neuronal morphology. Additionally, in the UMAP plot of snRNA-seq datasets from hippocampal neurons we found in each excitatory neurons subpopulation, the CTCF- cKO nuclei migrated closer into center. The study of Lipinski et al, 2020, present that elimination of both KAT3 genes leads to depletion in cell type specific markers and identity loss and therefore, KAT3- cKO nuclei migrated closer into center [23].

Regression in CTCF- cKO nuclei in every cluster of the excitatory neurons subpopulations, through differently expressed genes that related to regulation of cell population, suggest the crucial role of CTCF in maintaining of neuronal identity.

The current study determines the function of CTCF, follow changes in chromatin structure and gene transcription, in neuronal identity which influence behavioral changes. Therefore, CTCF has specific behavioral and neuronal functions in adulthood and is an important factor in neuronal maintenance.

**Figure 8-1 (supplementary): Classification of CamkIIa expressing cells to subgroups through specific cell-type markers.**

## Acknowledgements

We would like to thank Tal Katz Ezov, Nitsan Fourier and Maor Hatoel from the Technion Genomic Center for their assistance in library preparation, sequencing, quality control and initial bioinformatic analysis of the single nuclei sequencing. current study was funded by Israel Science Foundation Grant 898/17.

---

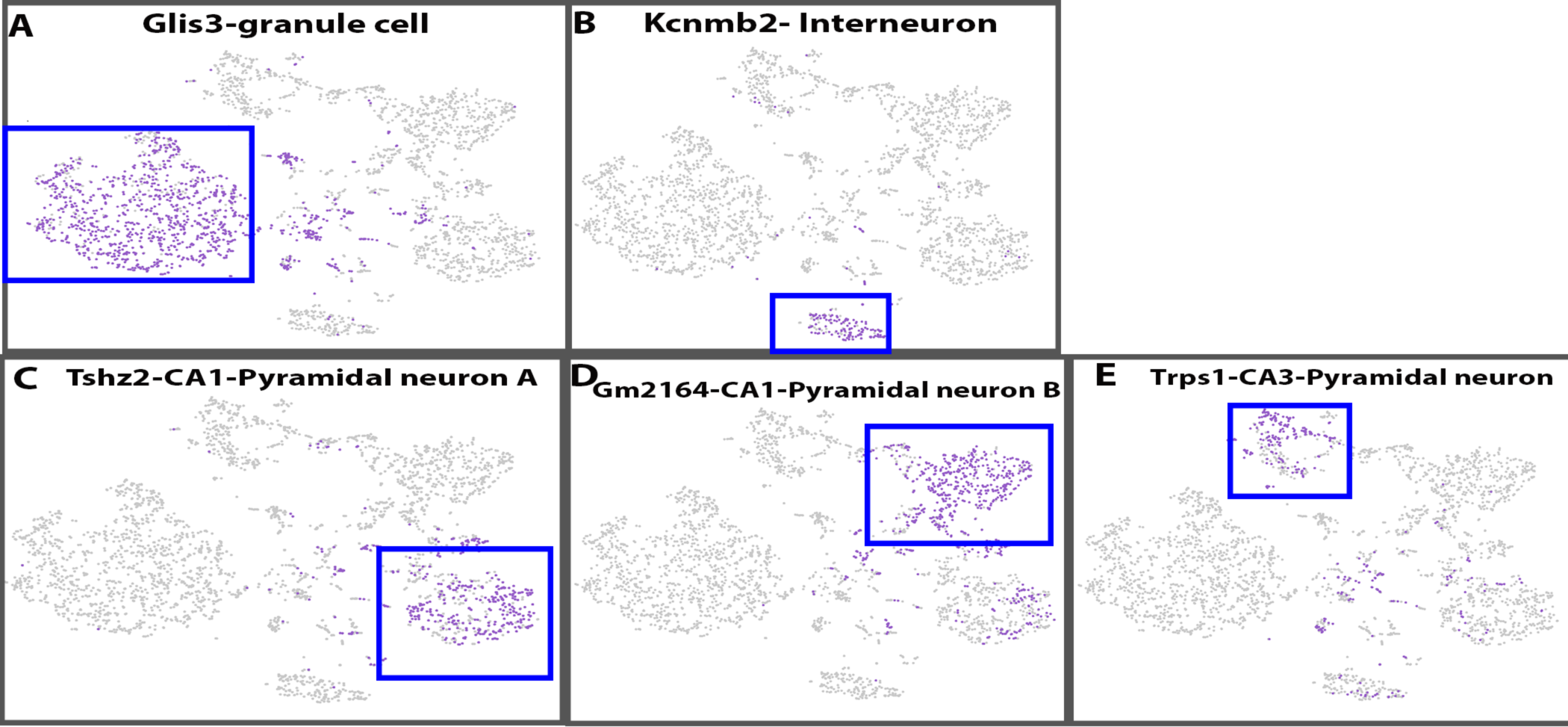

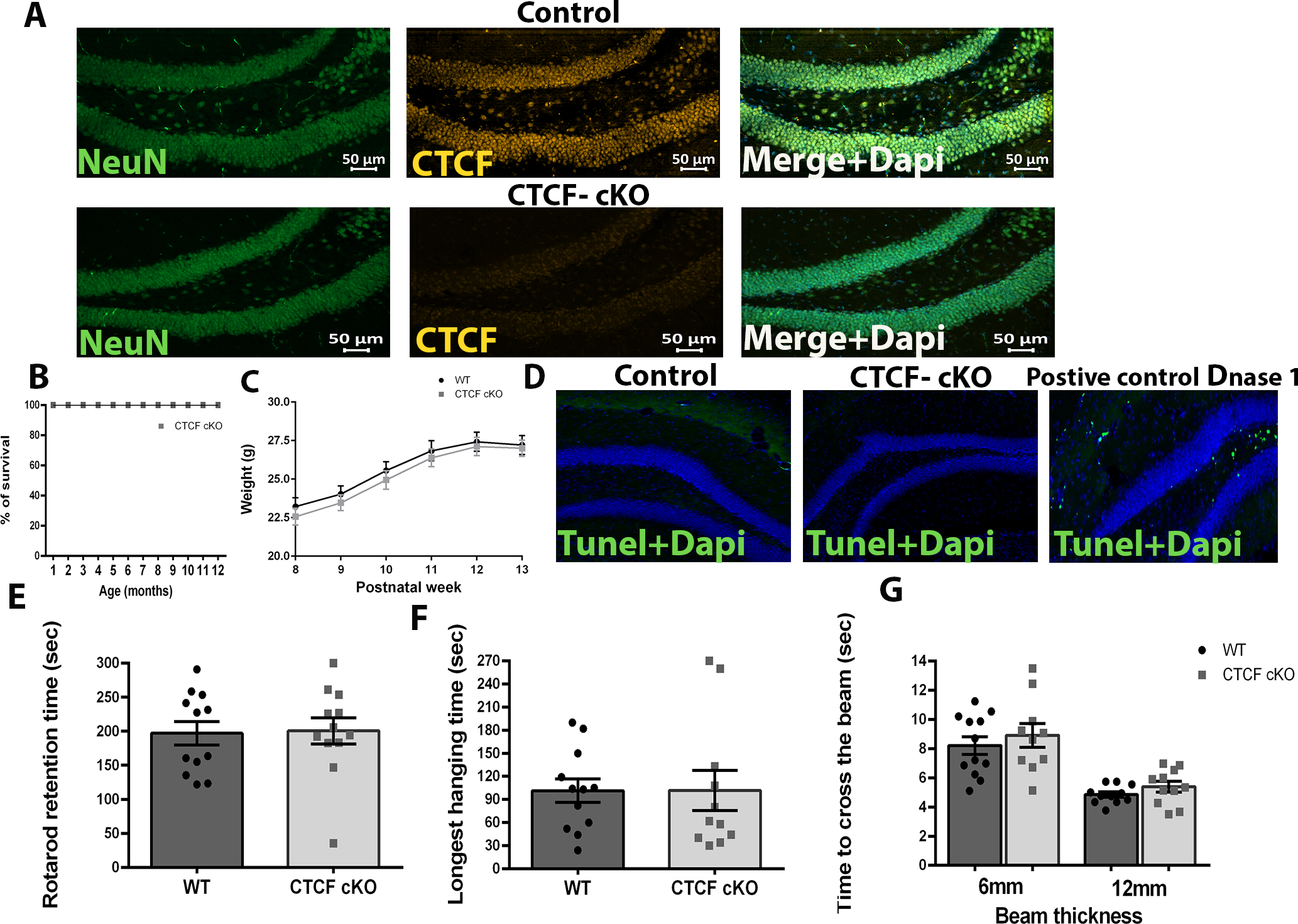

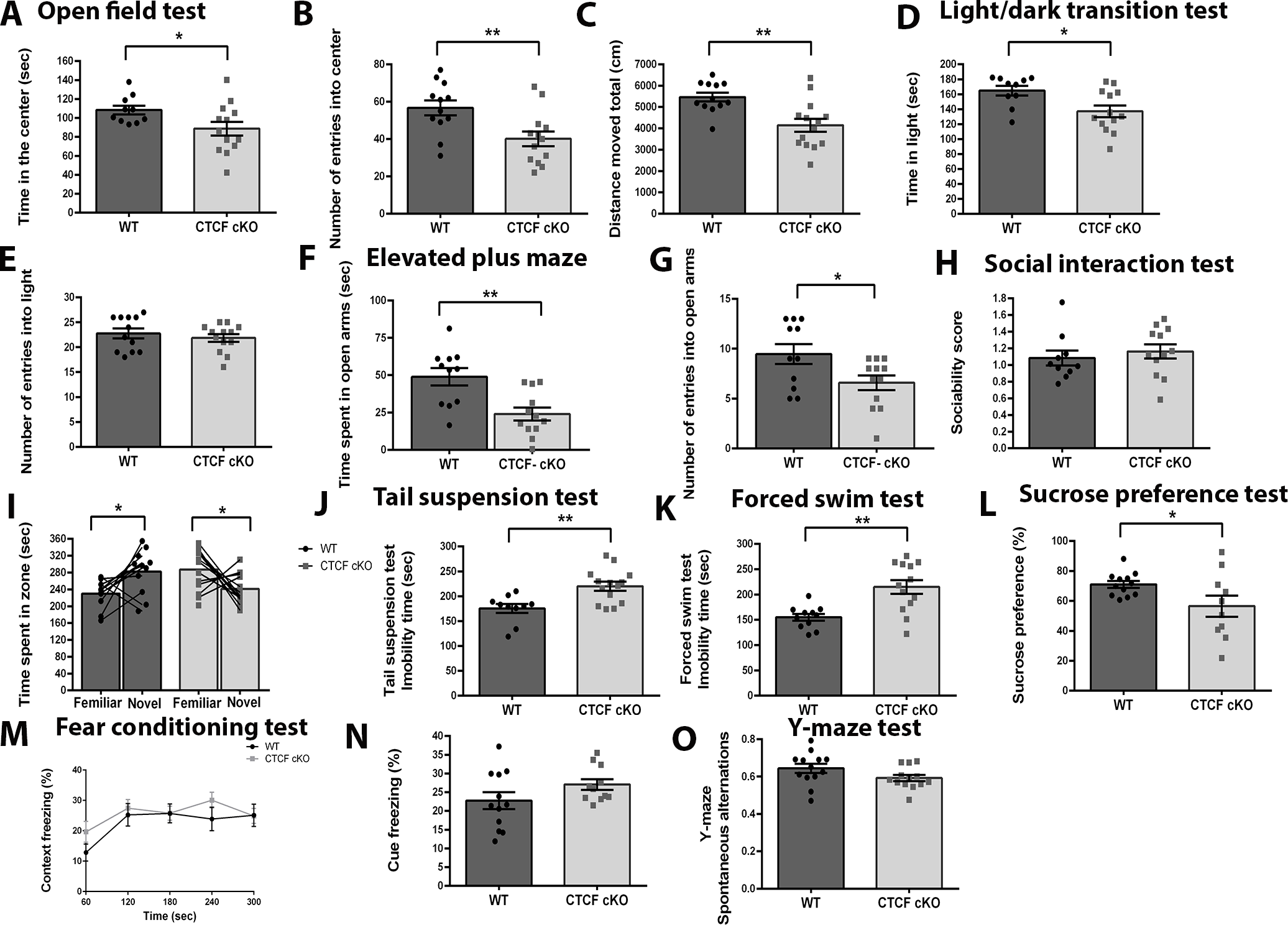

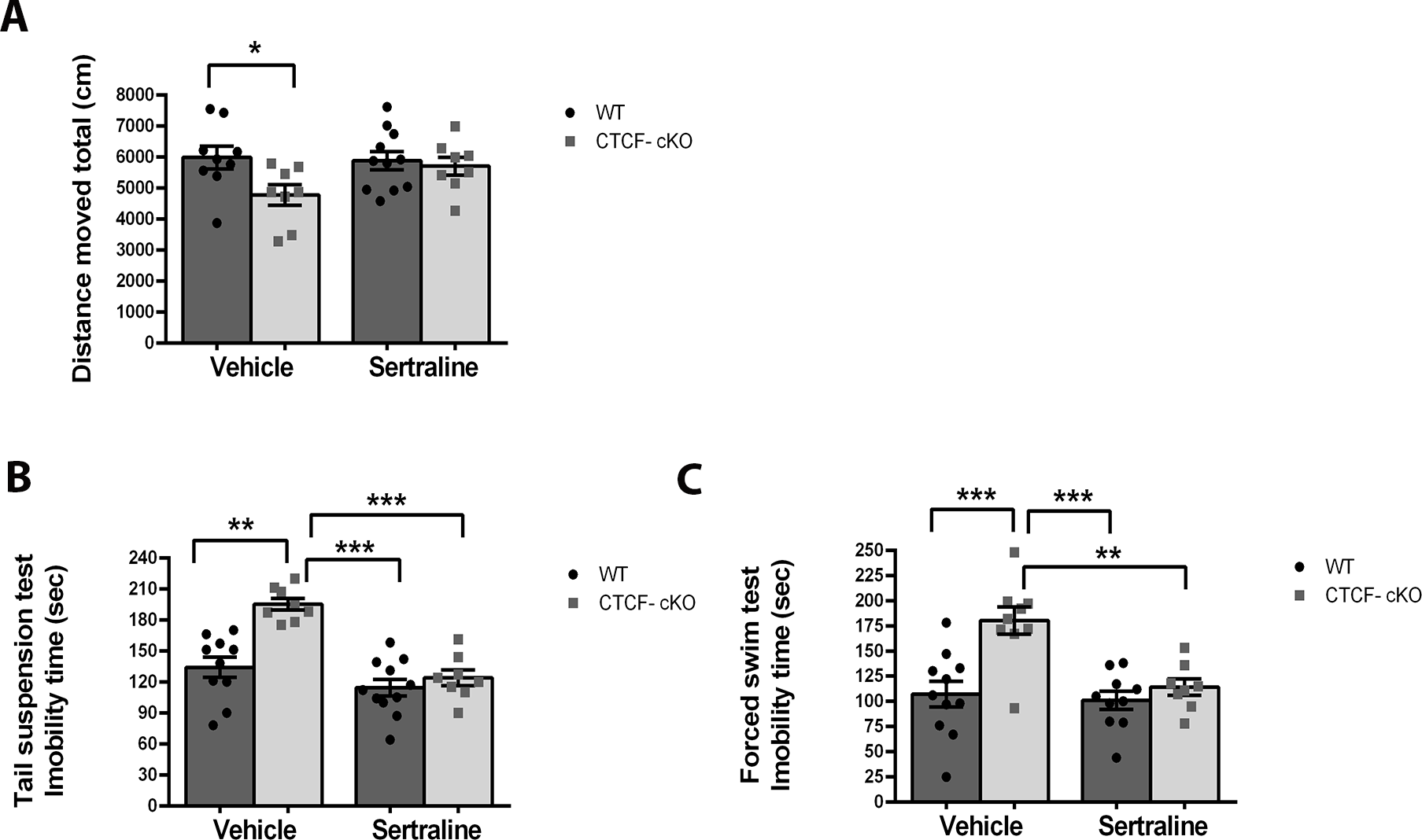

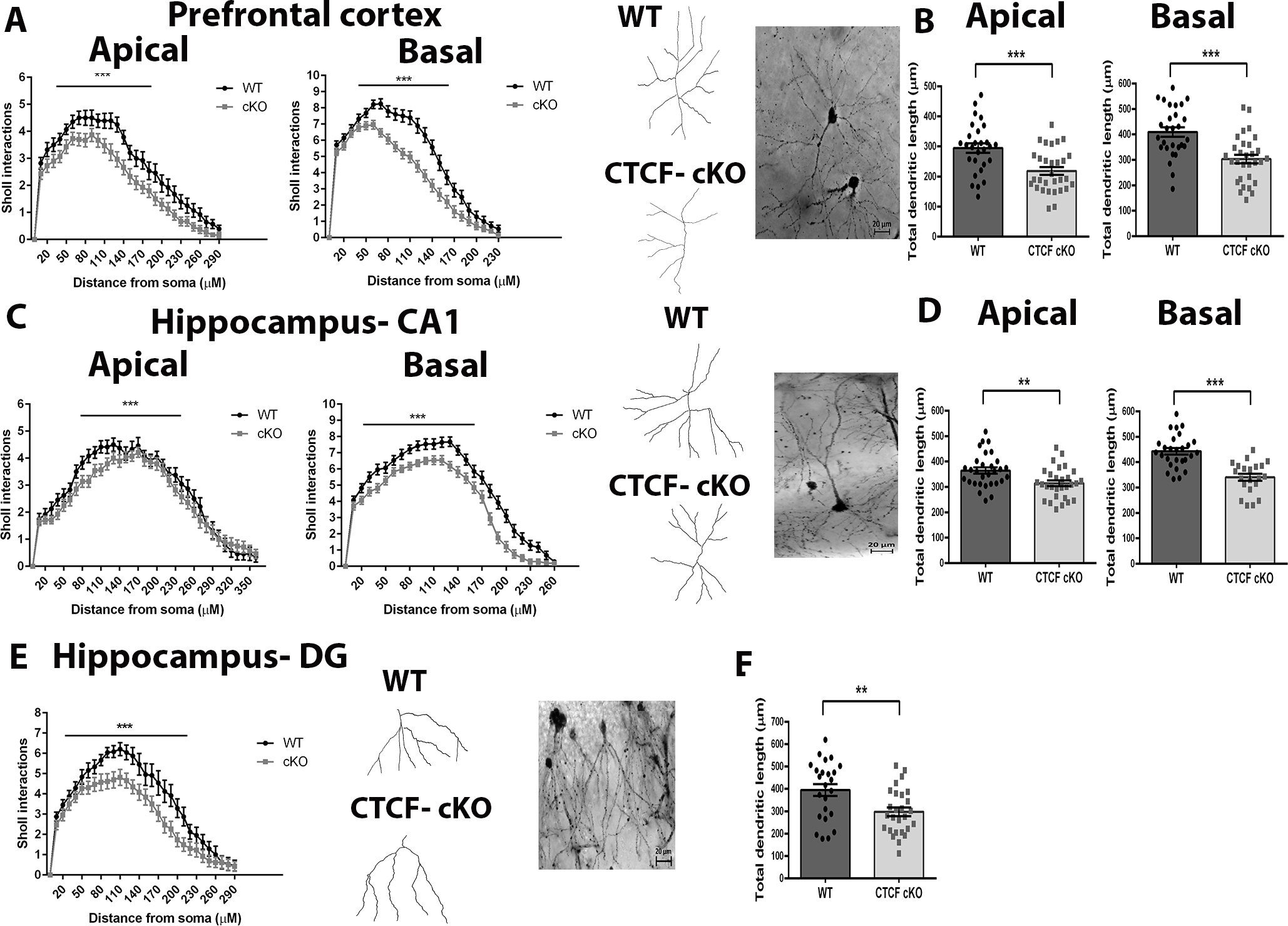

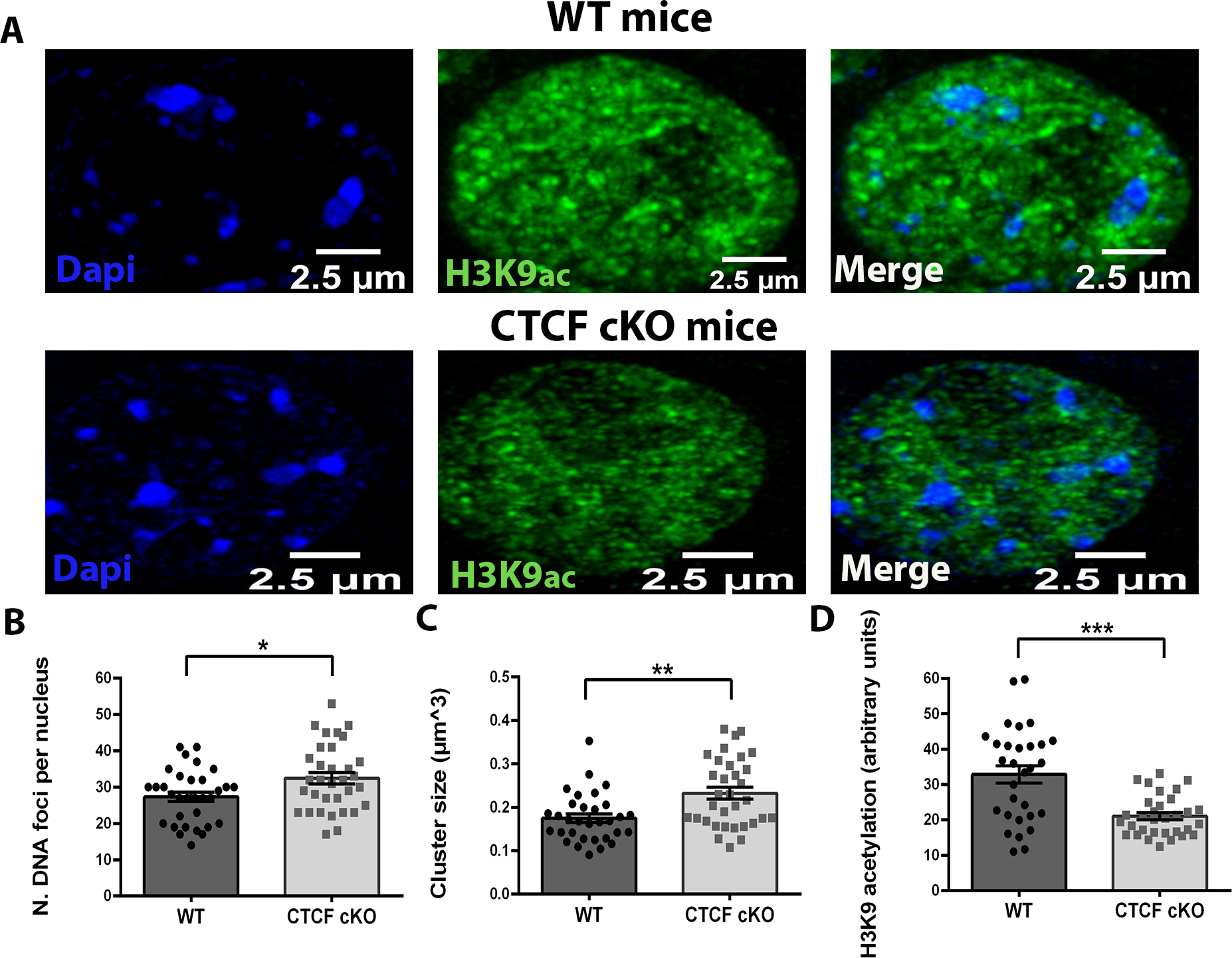

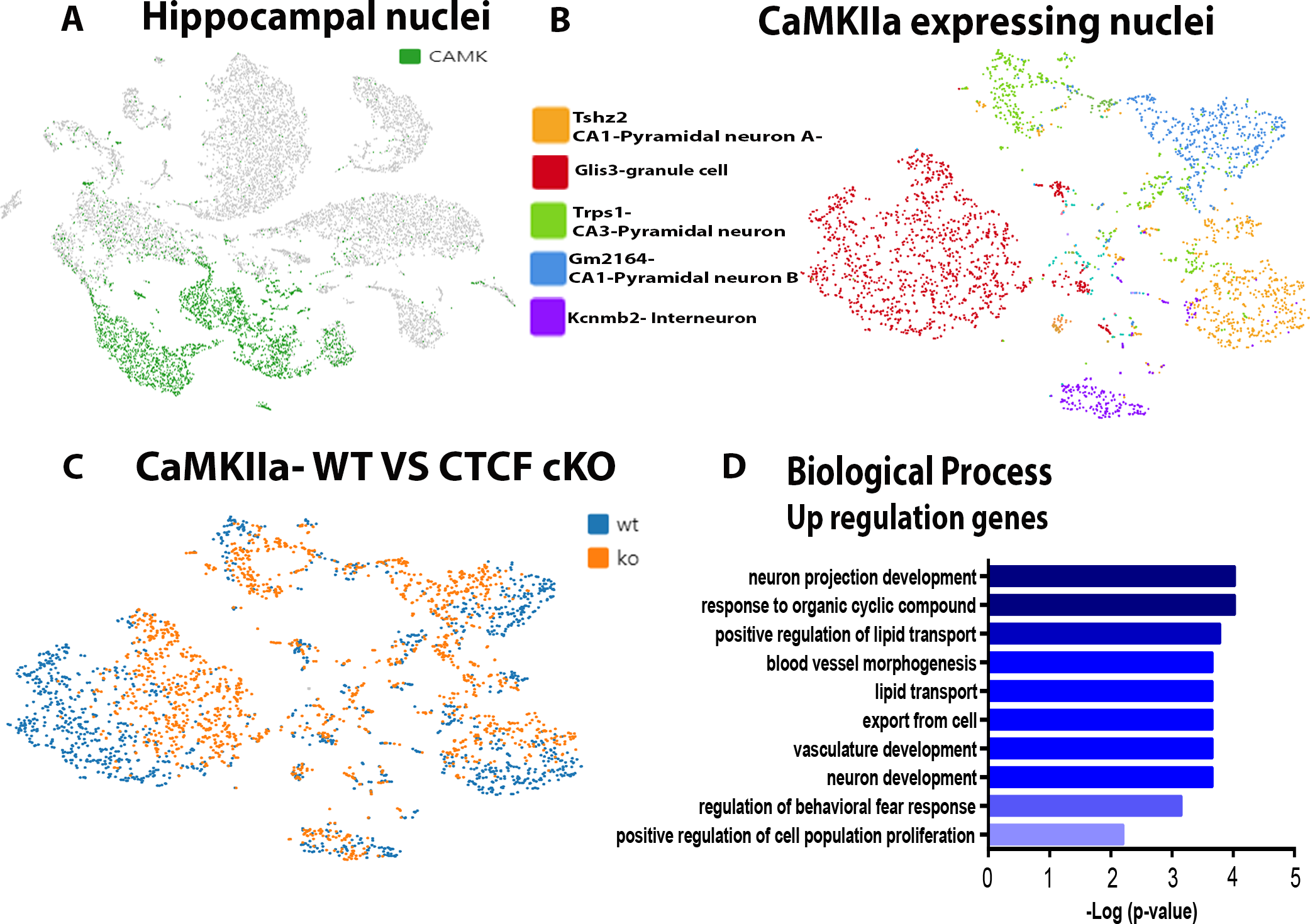

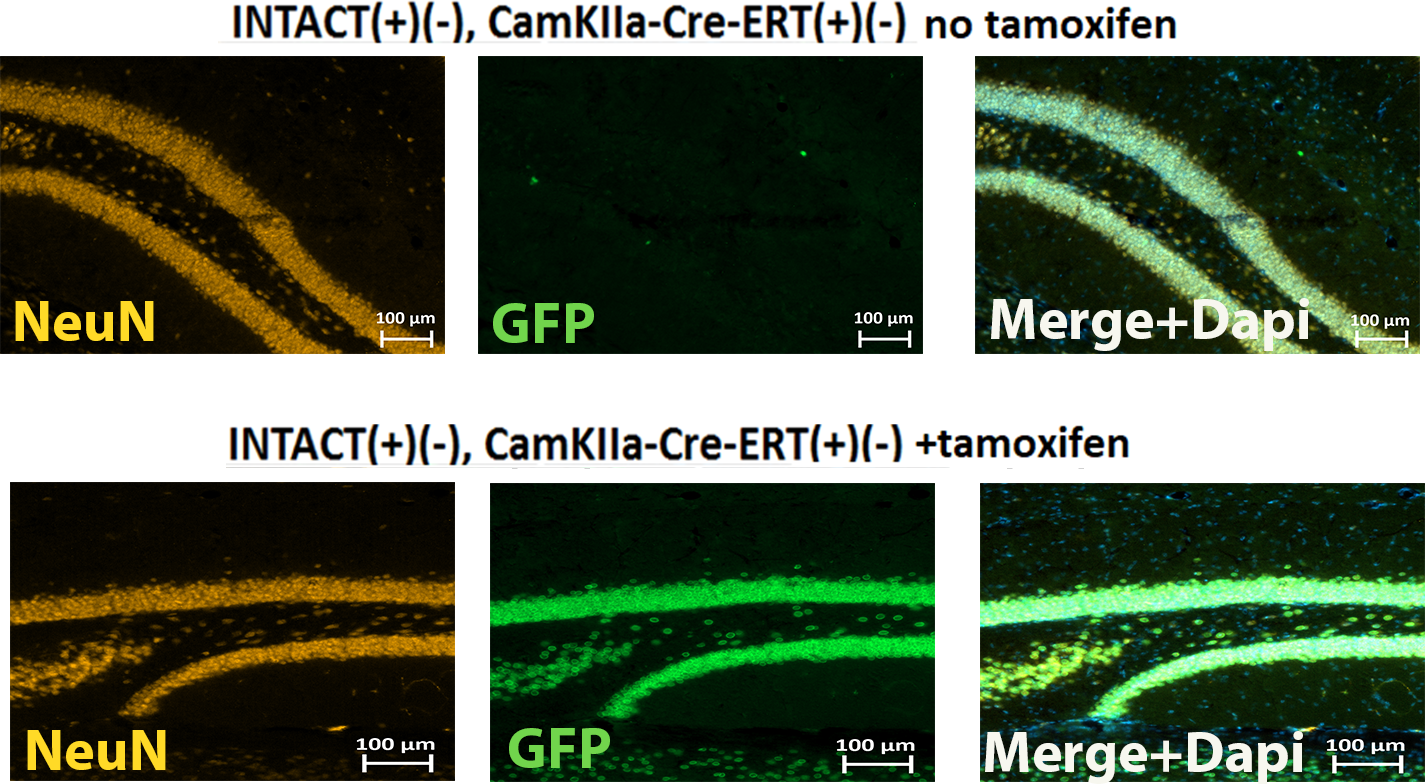

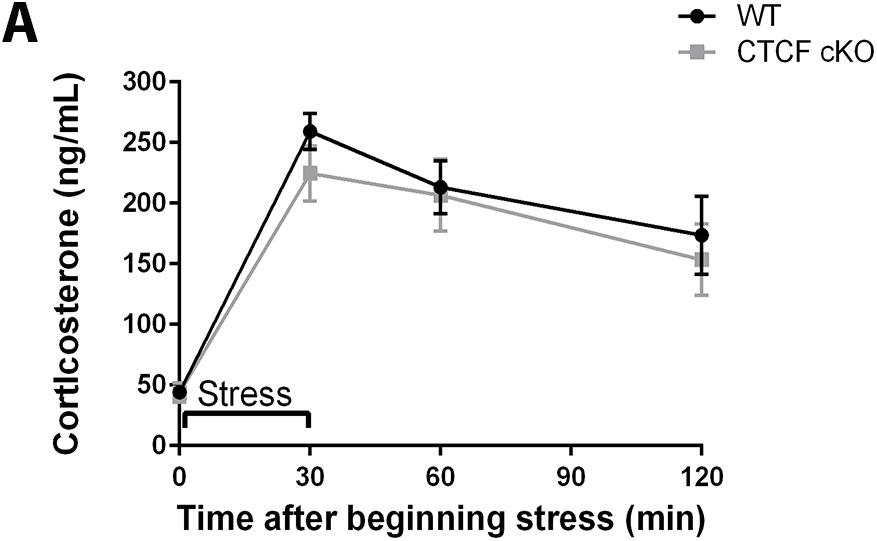

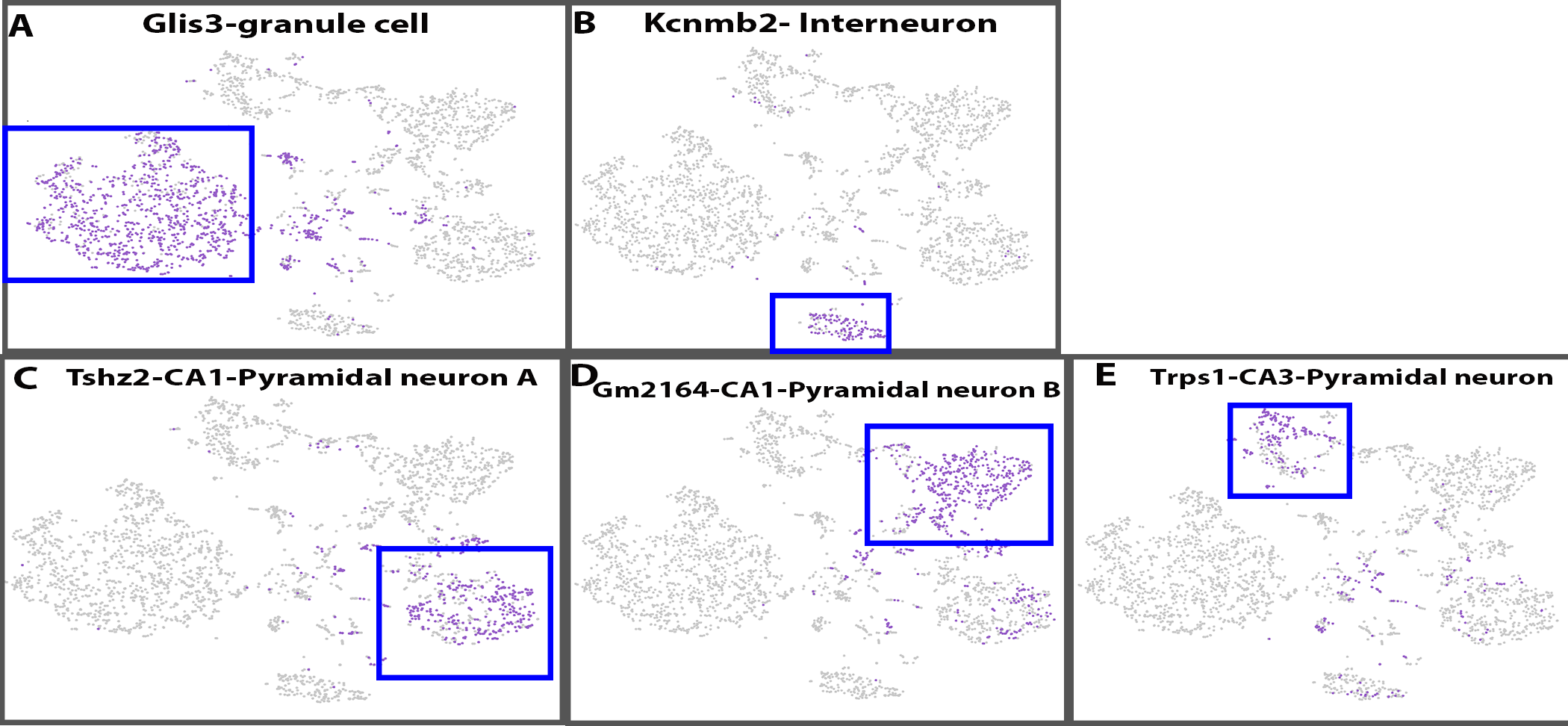

